# Rapid screening and identification of genes involved in bacterial extracellular membrane vesicle production using a curvature-sensing peptide

**DOI:** 10.1101/2024.05.20.594893

**Authors:** Hiromu Inoue, Kenichi Kawano, Jun Kawamoto, Takuya Ogawa, Tatsuo Kurihara

## Abstract

Bacteria secrete extracellular membrane vesicles (EMVs). Physiology and biotechnological applications of these lipid nanoparticles have been attracting significant attention. However, the molecular basis of EMV biogenesis has not yet been fully elucidated. In the previous research, an N-terminus-substituted FAAV peptide labeled with NBD (nFAAV5-NBD) was developed, which can sense a curvature of a lipid bilayer and selectively bind to EMVs in the presence of the cells. Herein, we applied nFAAV5-NBD to a screening of hyper- and hypo-vesiculation transposon mutants of a Gram-negative bacterium, *Shewanella vesiculosa* HM13, to identify the genes involved in EMV production. As a result, for 16 and six genes, we found that transposon insertion within or near the gene caused hyper- and hypo-vesiculation, respectively. Targeted gene-disrupted mutants of the identified genes demonstrated that the lack of putative dipeptidyl carboxypeptidase, glutamate synthase β-subunit, LapG protease, metallohydrolase, RNA polymerase sigma-54 factor, inactive transglutaminase, PepSY domain-containing protein, and Rhs-family protein caused EMV overproduction. On the other hand, disruption of the genes encoding putative phosphoenolpyruvate synthase, D-hexose-6-phosphate epimerase, NAD-specific glutamate dehydrogenase, and sensory box histidine kinase/response regulator decreased EMV production. This study shows the utility of a novel screening method using a curvature-sensing peptide and fundamental information on the genes related to EMV production.

**IMPORTANCE:** Conventional methods for isolation and quantification of EMVs are generally time-consuming. nFAAV5-NBD can detect EMVs in the culture without separating EMVs from cells. *In situ* detection of EMVs using this peptide facilitated screening of the genes related to EMV production. We succeeded in identifying a number of genes involved in the EMV production of *S. vesiculosa* HM13, and the strains obtained in this study would contribute to the elucidation of bacterial EMV formation mechanisms. Additionally, the hyper-vesiculating mutants obtained in this study would be useful in the application of this bacterium, such as in the production of useful substances as cargoes of EMVs and in the production of surface-engineered vesicles.

## INTRODUCTION

Production of extracellular membrane vesicles (EMVs), nano-sized lipid particles, is commonly observed across all three domains of life (1). Bacteria produce EMVs containing various biomolecules such as proteins, peptidoglycan, phospholipids, and nucleic acids (DNA and RNA) (2). EMVs are involved in diverse biological processes, including horizontal gene transfer, export of cellular metabolites, and intercellular communication (3). In addition to their importance in biological functions, EMVs have been used as a platform for heterologous protein production and as a carrier of biopharmaceuticals (4, 5).

Gram-negative bacteria produce EMVs by outer membrane blebbing and pinching off, by lytic release of the membrane fragments with bacteriolysis, and by corelease of outer membrane and inner membrane (2, 3). It has been documented that membrane blebbing results from a decrease in membrane integrity due to disruptions of crosslinking of the outer membrane and peptidoglycan (6, 7), local curvature formation by modification of asymmetric bilayer of the outer membrane (7–9), and membrane stress caused by the periplasmic accumulation of misfolded proteins, lipopolysaccharide (LPS), and peptidoglycan components (10–13). A variety of vesicle morphologies have been reported, including outer-inner membrane vesicles (14) and multilamellar and multivesicular outer membrane vesicles (15), which are formed by processes different from typical membrane blebbing and cell lysis. However, the molecular basis behind EMV biogenesis has not yet been fully clarified. Thus, comprehensive analysis of the genes related to EMV formation contributes to the elucidation of a novel EMV-producing mechanism.

To investigate EMV production-related genes, the selection of the mutants with altered EMV productivity from a gene knockout collection (7) or a random mutation library (6, 16) has been carried out. In these studies, EMV productivity of the mutants was evaluated by immunostaining using anti-LPS antibody (6, 7) or an antibody against a major cargo protein (16). As far as we know, no study has conducted a rapid evaluation of EMV productivity based on *in situ* detection of EMVs. In the previous research, a curvature-sensing peptide, FAAV, was designed based on the peptide sequence derived from sorting nexin protein 1 (SNX1), which is a Bin/Amphiphysin/Rvs (BAR) family protein (17). Then, by modifying the N-terminus and labeling with a fluorescence group, nitrobenzoxadiazole (NBD), N-terminus-substituted FAAV5 labeled with NBD (nFAAV5-NBD) was developed (18). In the presence of cells, nFAAV5-NBD can selectively bind to the lipid packing defects on EMVs, where the arrangement of the lipid bilayer is disordered due to the high membrane curvature. Therefore, we considered that this peptide could be applicable to the selection of mutants with changes in EMV productivity without removing the cells from the culture.

*Shewanella vesiculosa* HM13, a Gram-negative bacterium isolated from the intestine of a horse mackerel, produces larger amounts of EMVs with less variation in particle size than other well-known *Shewanella* species and *Escherichia coli* (19). Genetic approaches to clarify the molecular basis behind the EMV production of this strain are expected to be a clue for elucidating important mechanisms of bacterial EMV production and morphogenesis. EMVs secreted by this strain harbor a 49 kDa protein, named P49, of unknown function as their single major cargo protein (19, 20). Using P49 as a carrier to transport heterologous proteins to EMVs, this strain could be useful as a host for the production of EMVs with desired properties.

In this research, we established the rapid screening method for the genes related to EMV production using random transposon mutagenesis and nFAAV5-NBD. The screening resulted in the selection of 18 hyper- and eight hypo-vesiculating strains. Based on the identified transposon insertion sites, we conducted targeted gene disruption of individual genes for verification. Through the characterization of EMV production, cell proliferation, and protein secretion of the targeted gene-disrupted strains, eight and four genes whose disruption increased and decreased EMV productivity, respectively, were identified. In brief, the EMV productivity of *S. vesiculosa* HM13 was suggested to be regulated by protein quality control, extracellular environmental signal sensing, and glutamate metabolism. This study provides a novel screening method using a curvature-sensing peptide and fundamental information on genes related to EMV production.

## RESULTS

### Evaluation of EMV-selective binding of nFAAV5-NBD

To validate whether the NBD-labeled curvature-sensing peptide, nFAAV5-NBD, can exhibit a high ability to detect the EMV concentration in the culture, nFAAV5-NBD was added to the EMV-containing fractions (culture, cell-free supernatant, and EMVs) and the EMV-free fractions (cell suspension and post-vesicle fraction (PVF)). When the fractions without dilution were used, the culture and the cell-free supernatant showed about five-fold higher fluorescence intensity of NBD than the cell suspension (Fig. 1A). The fluorescence intensity of the EMV-containing fractions (culture, cell-free supernatant, and EMVs) increased with increasing relative concentration, whereas such a trend was not observed for the EMV-free fractions (cell suspension and PVF) (Fig. 1B). Thus, this peptide can be applied to the *in-situ* evaluation of EMV productivities of *S. vesiculosa* HM13.

**Fig. 1.**
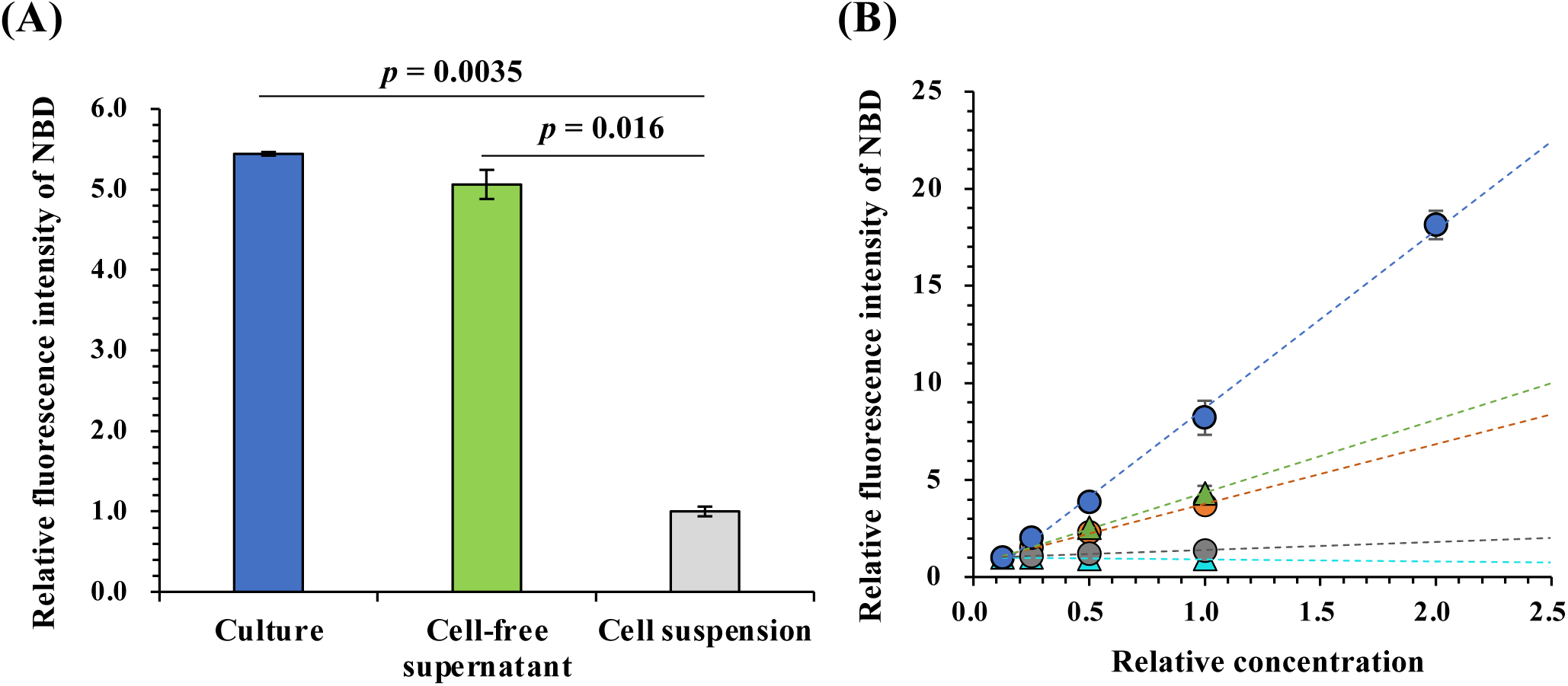
Detection of EMVs in EMV-containing fractions using nFAAV5-NBD. **(A) Detection of EMVs in culture and cell-free supernatant using the curvature-sensing peptide nFAAB5-NBD** Each undiluted fraction (relative concentration of 1.0) was incubated with nFAAV5-NBD. The data are the means +/− standard deviation of the values from three independent batches. *p* value (Student’s *t*-test). **(B) Measurement of concentration-dependent changes in NBD fluorescence intensity for each fraction** A relative concentration of 1.0 indicates a concentration of each component in the original culture. Culture (orange circle), cell-free supernatant (green triangle), PVF (cyan triangle), cell suspension (gray circle), and EMVs (blue circle) were incubated with nFAAV5-NBD, and the fluorescence intensity of NBD was quantified. The data are the means +/− standard deviation of the values from three independent batches.

### *In situ* rapid detection of EMV productivities of random transposon mutants (Tn mutants)

The randomness of transposon insertions into the genome was ensured by the identification of transposon insertion sites in ten randomly selected colonies from the library (data not shown). In the first selection, 10,100 Tn mutants were statically cultured in 96-well plates, and their EMV productivities were evaluated by NBD fluorescence intensity and OD value. The logarithm of fold change in EMV production by the mutants showed a unimodal broad distribution (average: 0.017, median: 0.020) (Fig. 2A), indicating that the EMV productivities of screening-subjected mutants were certainly variable. After the selection, 85 and 56 mutants were selected as hyper- and hypo-vesiculating candidates according to their fold changes of more than 2.0 and less than 0.5, respectively (Fig. 2B). The former and latter were in the top 0.84% and bottom 0.55% of the screening subjects.

**Fig. 2.**
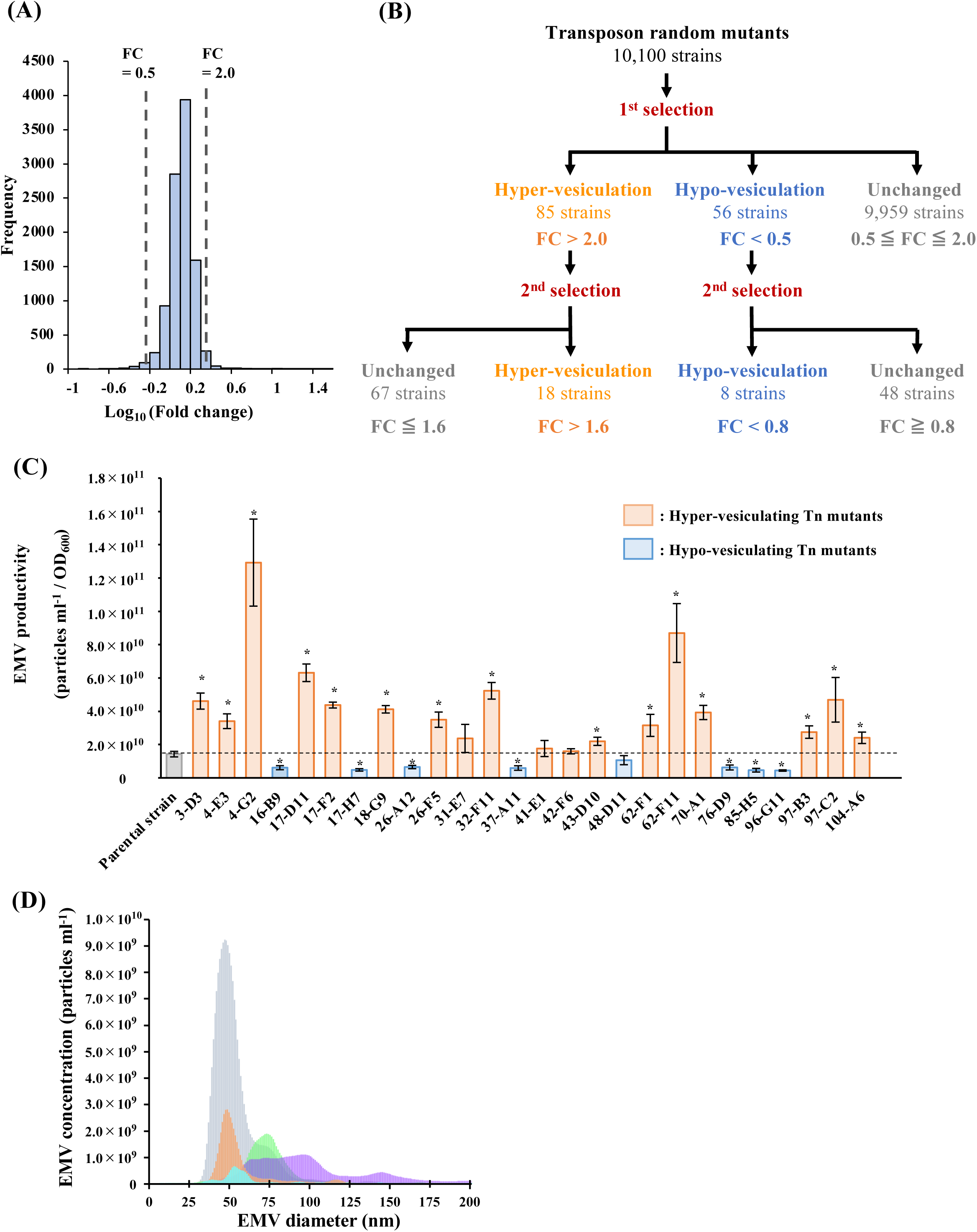
Selection of the Tn mutants using nFAAV5-NBD. **(A) Histogram indicating the distribution of fold change of EMV production in a library of random Tn mutants in the first selection** The mutants with a fold change of less than 0.5 and more than 2.0 (outside vertical dashed lines) were subjected to the second selection. **(B) Flowchart showing the number of subjects and thresholds for each selection** The number of mutants subjected to the present screening was 10,100, which was equivalent to 2.4 times the predicted number of the genes of strain HM13. **(C) EMV productivity of the parental strain and Tn mutants** EMV productivity of the strains was quantified by NTA. Average +/− SE; the parental strain, n = 10; Tn mutants, n = 3. ∗: *p* < 0.05 (Student’s *t*-test). **(D) Hydrodynamic diameter of EMVs of the parental strain and Tn mutants** Particle diameter was quantified by NTA. The parental strain, n = 5; Tn mutants, n = 3. Each graph shows the average. Green: the parental strain, cyan: strain 26-A12, orange: strain 26-F5, gray: strain 32-F11, purple: strain 70-A1.

To further select the strains whose EMV productivity increases or decreases under aerobic culture condition, 141 candidates selected from the first screening were aerobically cultivated and subjected to the evaluation of EMV productivity using nFAAV5-NBD as described in Materials and Methods. Eighteen and eight strains were selected as hyper- and hypo-vesiculating strains according to the fold change of more than 1.6 and less than 0.8, respectively (Fig. 2B).

### Characterization of selected Tn mutants

The EMV fractions of the selected Tn mutants were isolated, and the EMV concentration and hydrodynamic diameters were measured by nanoparticle-tracking analysis (NTA) (Fig. 2CD, Fig. S1A). When EMVs were quantified using nFAAV5-NBD and NTA, the hyper- and hypo-vesiculation phenotypes were consistently observed (Table S1), although there were four strains whose EMV productions showed no significant differences from that of the parental strain (*p* > 0.05) (Fig. 2C). Among eighteen mutants selected as hyper-vesiculating strains, fifteen mutants produced significantly larger amounts of EMVs compared to the parental strain. Strain 4-G2 exhibited the highest EMV productivity, and its vesiculation level was 9.0-fold higher than that of the parental strain (Fig. 2C). On the other hand, EMV productivity was significantly decreased in the hypo-vesiculating mutants except for strain 48-D11, and the lowest EMV productivity was observed for strain 96-G11, which produced 0.31-fold smaller amount of EMVs than the parental strain (Fig. 2C).

The hydrodynamic diameters of EMVs produced by the parental strain and the Tn mutants were compared. Analysis of particle size distribution by NTA indicated that the parental strain mainly produced EMVs with a diameter of 60–90 nm (Fig. 2D). Although most strains isolated as hyper- and hypo-vesiculation mutants showed the same particle size distribution as the parent strain (Fig. S1A), the hyper-vesiculating strains 26-F5 and 32-F11 and the hypo-vesiculating strain 26-A12 mainly produced smaller EMVs with a diameter of 40–60 nm than the parental strain (Fig. 2D). The hyper-vesiculating strain 70-A1 produced slightly larger EMVs than the parental strain with increased size variation (Fig. 2D).

Overall, we successfully selected the Tn mutants with increased or decreased EMV productivity using nFAAV5-NBD.

### Identification of transposon insertion sites

Transposon insertion sites were identified by inverse PCR (21) or single-primer PCR (22). Transposon insertion sites identified for eighteen hyper-vesiculating strains are listed in Table 1. In brief, transposons were inserted into the coding region of 16 genes and one 5’-untranslated region (UTR). Among the transposon-inserted genes, two genes were predicted to encode inner membrane proteins (HM2192 and HM502) containing predicted seven and eight transmembrane helices, respectively, two genes to encode outer membrane proteins (HM1685 and HM2827) not containing predicted transmembrane regions, and six genes to encode cytoplasmic proteins (HM3484, HM1858, HM2418, HM3946, HM2929, and HM3068). For the other six proteins (HM4090, HM1880, HM1399, HM3230, HM1963, and HM1528), PSORTb and SOSUI programs did not predict the localization or did not yield consistent results. The transposon was inserted in the 5’-UTR of *hm2720* and *hm2721*, encoding a putative Gate domain-containing spore maturation protein and 16S rRNA methyltransferase RsmF, respectively. In the 41-E1 and 70-A1 strains, the transposon was inserted into different positions of *hm2192*, encoding a putative inactive transglutaminase. By BLASTP search, there were no sequence homologs of HM1880, HM1858, and HM1528 with predictable functions. However, the AlphaFold2 predicted models of HM1880 and HM1528 showed structural similarity to LapG protease from *Legionella pneumophila* (Cα RMSD: 1.8 Å) and an adaptor protein required for acetylation of the alginate exopolysaccharide from *Pseudomonas aeruginosa* (Cα RMSD: 2.1 Å), respectively (Fig. S2). There was no protein with a similar three-dimensional structure to HM1858.

**Table 1.**
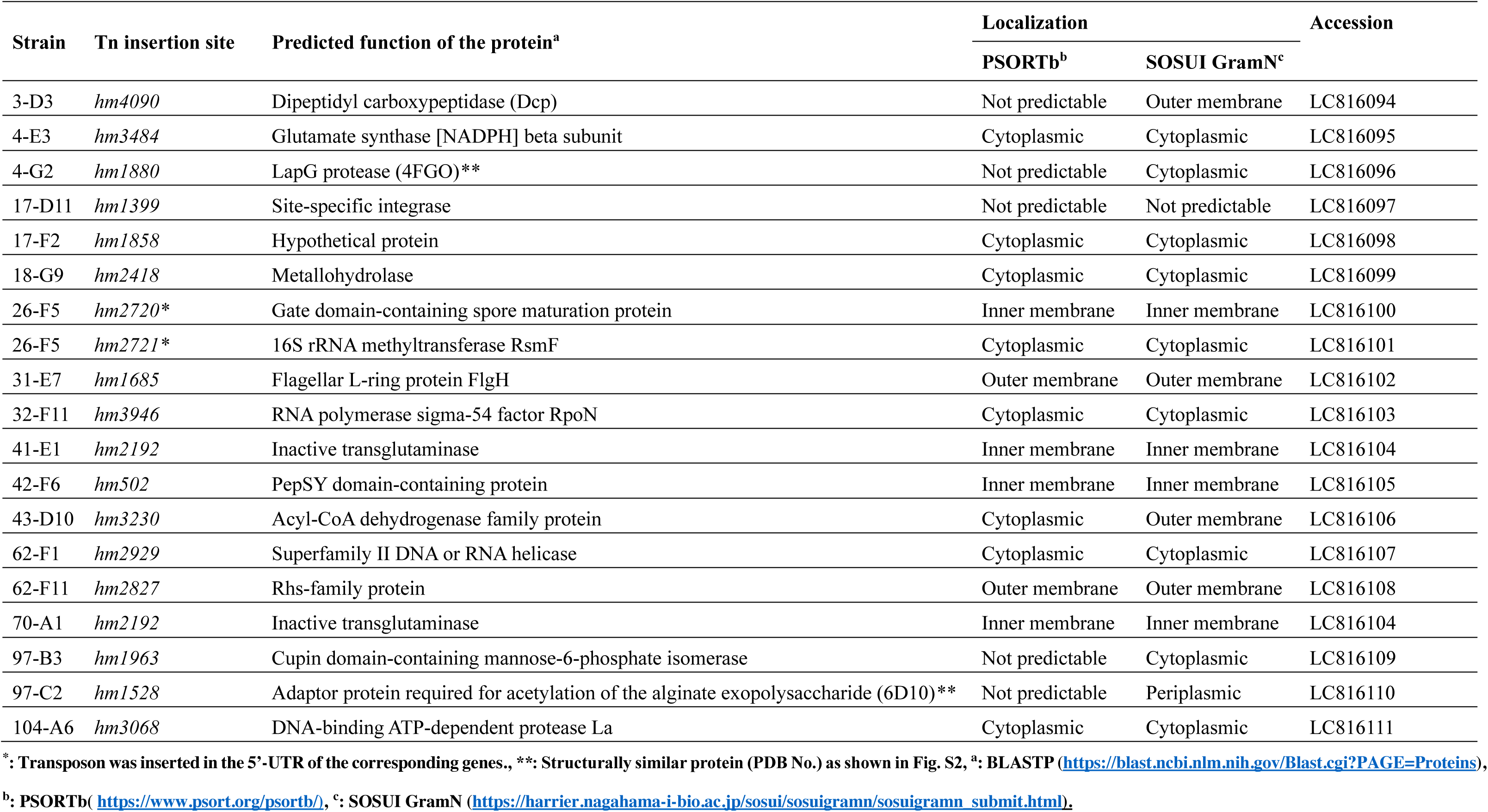
Tn insertion sites of the hyper-vesiculating strains.

On the other hand, in hypo-vesiculating strains, transposons were inserted into six genes and one 5’-UTR (Table 2). Among the genes with transposon insertions, one gene was predicted to encode inner membrane protein (HM3454) containing predicted seven transmembrane helices, and two genes were to encode cytoplasmic proteins (HM2766 and HM369). For the other proteins (HM2775, HM2704, and HM3986), PSORTb and SOSUI programs did not yield consistent results. In the 37-A11 strain, the transposon was inserted into 5’-UTR of *hm238* and *hm239*, encoding a hypothetical protein and 23S rRNA methylase RlmKI, respectively. The function of HM238, a short peptide chain consisting of 39 amino acid residues, could not be predicted based on its sequence and predicted structure. In the 76-D9 and 96-G11 strains, the transposon was inserted into different sites of *hm2704*, encoding putative NAD-specific glutamate dehydrogenase.

**Table 2.** Tn insertion sites of the hypo-vesiculating strains.

| Strain | Tn insertion site | Predicted function of the protein<sup>a</sup> | Localization |  | Accession |
| --- | --- | --- | --- | --- | --- |
|  |  |  | PSORTb<sup>b</sup> | SOSUI GramN<sup>c</sup> |  |
| 16-B9 | *hm2766* | Phosphoenolpyruvate synthase | Cytoplasmic | Cytoplasmic | LC816112 |
| 17-H7 | *hm2775* | D-Hexose-6-phosphate epimerase | Not predictable | Extracellular | LC816113 |
| 26-A12 | *hm369* | Molybdopterin molybdotransferase MoeA | Cytoplasmic | Cytoplasmic | LC816114 |
| 37-A11 | *hm238\** | Hypothetical protein | Inner membrane | Not predictable | LC816115 |
| 37-A11 | *hm239\** | 23S rRNA methylase RlmKI | Cytoplasmic | Cytoplasmic | LC816116 |
| 48-D11 | *hm3454* | CcsA-related protein | Inner membrane | Inner membrane | LC816117 |
| 76-D9 | *hm2704* | NAD-specific glutamate dehydrogenase | Not predictable | Cytoplasmic | LC816118 |
| 85-H5 | *hm3986* | Sensory box histidine kinase/response regulator | Inner membrane | Cytoplasmic | LC816119 |
| 96-G11 | *hm2704* | NAD-specific glutamate dehydrogenase | Not predictable | Cytoplasmic | LC816118 |

\*: Transposon was inserted in the 5'-UTR of the corresponding genes., <sup>a</sup>: BLASTP (https://blast.ncbi.nlm.nih.gov/Blast.cgi?PAGE=Proteins),
<sup>b</sup>: PSORTb( https://www.psort.org/psortb/), <sup>c</sup>: SOSUI GramN (https://harrier.nagahama-i-bio.ac.jp/sosui/sosuigramn/sosuigramn_submit.html).

### Verification of changes in EMV production by targeted gene disruption

To examine whether the identified genes are involved in EMV production by *S. vesiculosa* HM13, we generated the targeted gene-disrupted mutants by single homologous recombination using a knock-out plasmid, pKNOCK-Km (23). Quantification of EMVs by NTA revealed that Δ*hm4090*, Δ*hm3484*, Δ*hm1880*, Δ*hm2418*, Δ*hm3946*, Δ*hm2192*, Δ*hm502*, and Δ*hm2827* had significantly higher EMV productivity, and their EMV productions were 2.5-, 3.1-, 4.9-, 2.2-, 3.6-, 1.9-, 2.6-, and 1.8-fold higher than that of the parental strain, respectively (Fig. 3A). Δ*hm2766*, Δ*hm2775*, Δ*hm2704*, and Δ*hm3986* exhibited 0.38-, 0.54-, 0.42-, and 0.44-fold lower EMV productions than the parental strain with significant differences (Fig. 3B). The genes disrupted in these strains are considered to be involved in EMV production by strain HM13.

**Fig. 3.**
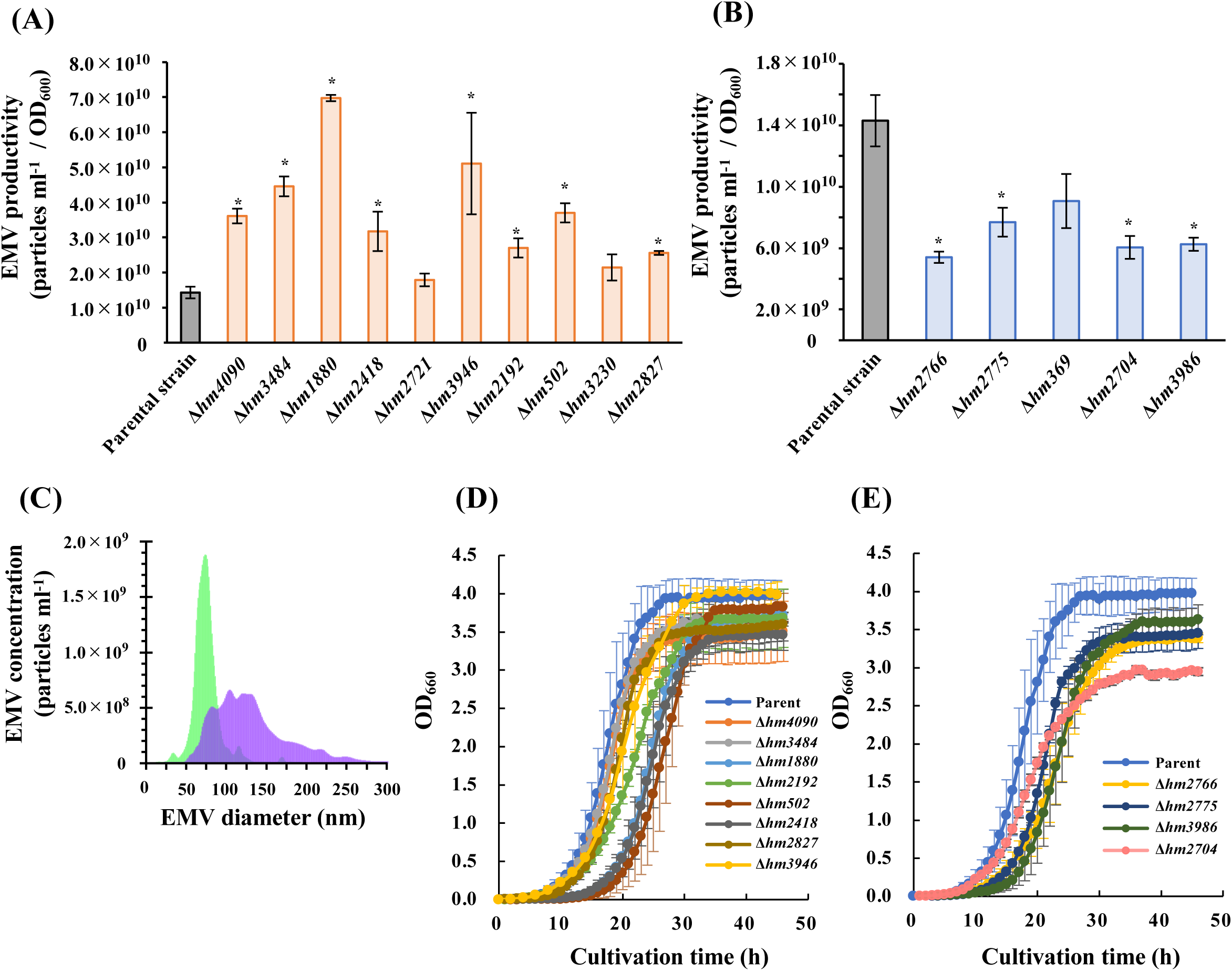
Characteristics of EMV production and growth of the gene-disrupted strains. **(A and B) EMV productivity of the parental strain and the targeted gene-disrupted strains (A: Hyper-vesiculating mutants, B: Hypo-vesiculating mutants)** EMV concentration was quantified by NTA. Average +/− SE, the parental strain, n = 10; the mutants, n = 3. ∗: *p* < 0.05 (Student’s *t*-test). **(C) Hydrodynamic diameter of EMVs of the parental strain and *hm2192*-disrupted strain (Δ*hm2192*)** Particle diameter was quantified by NTA. The parental strain, n = 5; Δ*hm2192*, n = 3. Each graph shows the average. Green: the parental strain, purple: Δ*hm2192*. **(D and E) Growth curve of the parental strain and the targeted gene-disrupted strains (D: Hyper-vesiculating mutants, E: Hypo-vesiculating mutants)** The growth of the strains was evaluated by monitoring OD660 with BioPhoto recorder TVS062CA. The parental strain, n = 5; the mutants, n = 3.

The hyper-vesiculating strain Δ*hm2192* produced EMVs with a large variation in size and mainly produced EMVs with a larger diameter of 75–125 nm than the parental strain (Fig. 3C). This morphological characteristic was consistent with that observed for the Tn mutant (70-A1) (Fig. 2D). This result suggests that HM2192 is related to the EMV particle size regulation in strain HM13. The other hyper-vesiculating strains (Fig. S1B) and hypo-vesiculating strains (Fig. S1C) mainly produced EMVs with size distributions almost identical to that of the parental strain.

### Growth characteristics of the targeted gene-disrupted strains

The growth of the parental strain and the targeted gene-disrupted strains was monitored (Fig. 3DE). Δ*hm1880*, Δ*hm2418*, Δ*hm502*, Δ*hm2766*, Δ*hm2775*, and *Δhm3986* showed longer lag phases than the parental strain. The doubling time of the parental strain was 2.2 h. Δ*hm2192* and Δ*hm2704* required longer proliferation times than the parental strain, with a doubling time of approximately 3.0−3.5 h. Although the defects in several identified genes caused prolonged lag phase and changes in doubling time and cell yields, these mutants did not exhibit cell aggregation and lysis until the early stationary phase under the tested conditions.

### Secretory protein production of the targeted gene-disrupted strains

The EMV fractions and PVFs collected from the parental strain and the targeted gene-disrupted strains were subjected to SDS-PAGE to analyze secretory protein productions (Fig. 4). As reported in our previous research (19, 20), a single major cargo protein P49 was observed in the EMV fraction collected from the parental strain with an apparent molecular mass of about 49 kDa. Similar to the parental strain, the bands presumed to be P49 were observed in all the EMV fractions collected from the mutants. However, many other protein bands were also observed in EMV fractions and PVFs from Δ*hm3484*, Δ*hm1880*, Δ*hm3946,* Δ*hm2192,* and Δ*hm2766*, indicating that these mutants secreted numerous proteins, other than P49, that were contained or not contained in EMVs.

**Fig. 4.**
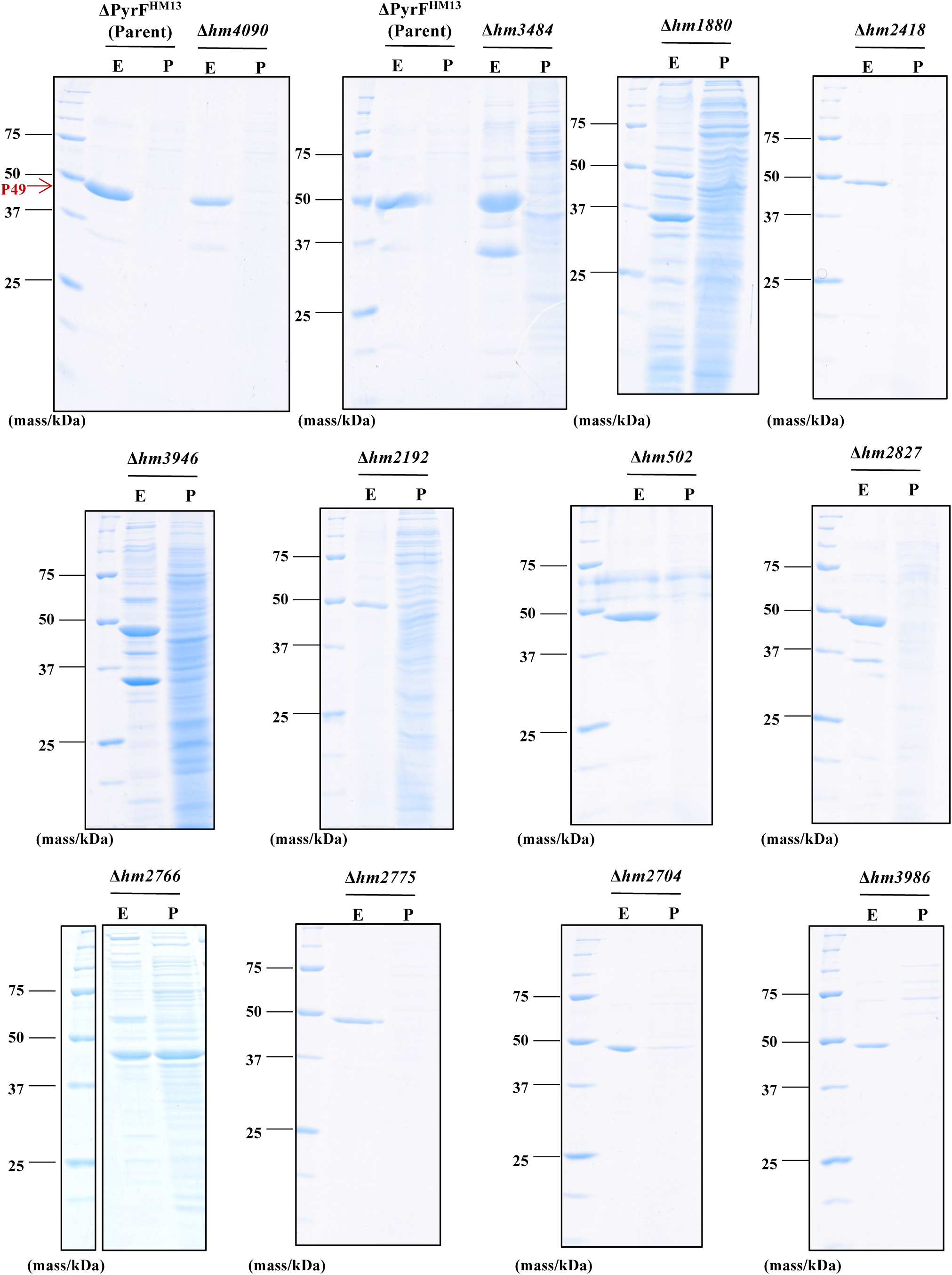
Analysis of the secretory proteins from the parental strain and the targeted gene-disrupted strains. EMVs and PVF from 300 µL of culture were subjected to TCA precipitation and SDS-PAGE. The gel was stained with Coomassie Brilliant Blue G-250. E: EMVs, P: PVF.

## DISCUSSION

### Summary of in situ rapid screening using a curvature-sensing peptide

The physiology and biotechnological applications of EMVs have been attracting significant attention (2, 3). However, the mechanism of bacterial EMV production has not been fully understood (24). Therefore, in this work, to identify the genes involved in EMV biogenesis by *S. vesiculosa* HM13 (19), we established and utilized a novel screening method combining random transposon mutagenesis with a curvature-sensing peptide, nFAAV5-NBD (18). The results of the binding assay to EMVs produced by strain HM13 indicated that nFAAV5-NBD could be applied to the *in situ* selection of the strains with variable EMV productivity (Fig. 1). By evaluating the amount of EMVs *in situ*, without separation of cells and EMVs, we succeeded in selecting mutants with altered EMV production (Fig. 2).

Expecting the genome-wide assessment, we subjected 10,100 mutants, which corresponds to approximately 2.4 times the estimated number of the genes in strain HM13, to the present experiment. Some of the genes were identified for more than one mutants in the screening (Tables 1 and 2). In the previous study, Kulp et al. conducted a high-throughput selection of hyper- and hypo-vesiculating strains from the whole genome knockout library of *Escherichia coli* mutant strains (Keio collection) (7). They elucidated that the knockout of the genes responsible for LPS and enterobacterial common antigen leads to hyper-vesiculation, whereas the knockout of the genes functioning in oxidative stress response pathways leads to hypo-vesiculation. However, the homologs of these genes were not identified in our present study. The present screening employed static culture in 96-well plates for the first selection of mutants (Fig. 2B), while in the screening conducted by Kulp et al., shaking culture was used (7), suggesting that these differences in culture conditions caused differences in the identified genes.

By nFAAV5-based screening, we selected eighteen hyper-vesiculating mutants and eight hypo-vesiculating mutants. We succeeded in identifying twenty-two genes and two 5’-UTRs where transposon was inserted (Tables 1 and 2). Notably, we elucidated the involvement of various proteins predicted to be localized at the outer membrane (HM1685 and HM2827) and the inner membrane (HM2192, HM502, and HM3454). Interestingly, disruption of *hm2192,* annotated as the gene encoding an inactive transglutaminase fused to seven transmembrane helices (7-TM) (Table S2), affected not only EMV production but also EMV size uniformity (Fig. 3A and 3C). Amidoligases including transglutaminase were reported to be involved in amide modifications to polysaccharides and proteins (25), but their function in EMV production remains unknown. Further analysis to elucidate the physiological function of this protein is needed.

### Enhancement of EMV production by defects of proteins involved in protein quality control

A gene encoding a homolog of dipeptidyl carboxypeptidase (Dcp) (HM4090) was identified from the hyper-vesiculating mutants. HM4090 was predicted to be a lipoprotein with an N-terminal lipid binding site by PSORTb program. Dcp is the protease that removes dipeptides from the C-termini of N-blocked tripeptides, tetrapeptides, and larger peptides (26). Because Dcp has broad substrate specificity (27), the Dcp homolog, HM4090, was speculated to perform a function of protein quality control in the outer membrane or periplasm. Membrane stress caused by an accumulation of misfolded proteins in the periplasm is one of the most renowned vesiculation mechanisms (2, 28). Previously, the deletion of DegP, a periplasmic protease/chaperon, increased EMV production by *E. coli* (28) without reducing the level of cross-link between the outer membrane and peptidoglycan (12). Therefore, membrane stress resulting from protein misfolding would cause hyper-vesiculation in Δ*hm4090*.

Δ*hm1880* exhibited 4.9-fold higher EMV production compared to the parental strain, and that was the highest EMV productivity in this study. Structure prediction by AlphaFold2 revealed that HM1880 has a LapG protease-like fold (Fig. S2AB). LapG is a periplasmic protease constituting the LapADG system, which is responsible for periplasmic proteolysis controlled by c-di-GMP-mediated signaling (29). In the c-di-GMP-rich condition, LapD, a transmembrane receptor, triggers sequestration of the protease LapG, thus preventing cleavage of the surface adhesin LapA (29, 30). In strain HM13, downstream of *hm1880*, the gene encoding HM1879, a LapD-like protein with EAL and GGDEF domains, is localized. However, the gene encoding a LapA homolog does not constitute this gene cluster. HM750, a homolog of T1SS-secreted agglutinin RTX showing 35.5% sequence similarity to LapA from *Pseudomonas fluorescens* (31), is included in an operon encoding T1SS, which is distant from the operon containing HM1880. Interestingly, although the involvement of the LapADG system in EMV production has not been elucidated, a LapADG-like system was identified as a regulator of biofilm formation in *S. oneidensis*, which is phylogenetically closely related to *S. vesiculosa* (32).

### Regulation of EMV production by signal transduction protein

HM3986, a sensory box histidine kinase/response regulator, contains a sensory box PAS domain and signal transduction domain BaeS (Table S2). PAS domains are conserved in three domains of life (33) and recognize the multiple stimuli from light (34), oxygen (35), and redox potential (36). In *E. coli*, the transposon insertion in *baeS* increased the expression of Spy (spheroplast protein Y), acting as a periplasmic ATP-independent chaperone with a function of protein quality control (37, 38). HM626 from *S. vesiculosa* HM13 showed 32% sequence homology to the spy protein from *E. coli* K12.

These findings suggest that *S. vesiculosa* HM13 has PAS and BaeS domain-mediated regulatory system for EMV production, and HM3986 might play a crucial role in the regulation of the accumulation of misfolded proteins in the periplasm by controlling the expression level of HM626, a homolog of spy protein.

### Involvement of metabolic enzymes in the EMV production

We revealed that disruption of two genes responsible for glutamate metabolism, *hm3484* and *hm2704*, results in increase and decrease of EMV production by strain HM13, respectively (Fig. 3AB). Glutamate metabolism is strictly regulated by GS/GOGAT and GDH cycles that comprise glutamate synthase and glutamate dehydrogenase (39). Disruption of the genes coding for a glutamate synthase β-subunit homolog, HM3484, or a glutamate dehydrogenase homolog, HM2704, would affect the cellular glutamate level. Glutamate is a major precursor of glutathione (40) and peptidoglycan (41), functioning in the scavenging of reactive oxidative species (ROS) and in membrane stability, respectively. Abundant ROS in the cell generates membrane stress and induces EMV production (42). These results suggest that EMV production by *S. vesiulosa* HM13 is regulated by the intracellular glutamate levels and its derivatives.

### Transposon insertion into non-coding regions

In Tn mutants 26-F5 and 37-A11, the transposon insertion site was identified in 5’-UTRs. The insertion of an exogenous sequence in 5’-UTR would cause polar effects on downstream genes (43). In mutant 26-F5, the transposon was inserted into the 5’-UTR of *hm2720* and *hm2721*, disrupting the putative promoter region of *hm2721* and separating the putative promoter of *hm2720* from the coding region (Fig. S3A). The transposon was inserted into the region between the putative promoter of *hm239* and its coding region, probably without affecting the expression of *hm238*, in mutant 37-A11 (Fig. S3B). Therefore, the genes encoding putative Gate domain-containing spore maturation protein (HM2720), 16S rRNA methyltransferase RsmF (HM2721), and 23S rRNA methylase RlmKI (HM239) possibly play important roles in suppression or facilitation of EMV production of *S. vesiculosa* HM13.

### Cell proliferation and protein secretion

Further analysis of the mutants revealed that four hypo-vesiculating mutants, Δ*hm2766*, Δ*hm2775*, Δ*hm3986*, and Δ*hm2704*, and four hyper-vesiculating mutants, Δ*hm1880*, Δ*hm2192*, Δ*hm502*, and Δ*hm2418*, exhibited growth retardation (Fig. 3DE). Among these mutants, except for Δ*hm2192 and* Δ*hm2704*, the lag phase was prolonged compared to that of the parental strain. In bacterial growth, the lag phase is a preparative phase of nonreplication for the subsequent logarithmic growth phase. During the lag phase, repair and replacement of damaged intracellular components proceed along with metabolic reorganization (44). The SDS-PAGE analysis showed that PVFs of Δ*hm3484*, Δ*hm1880*, Δ*hm3946*, Δ*hm2192*, and Δ*hm2766* contained a variety of protein bands not detected in the parent strain (Fig. 4). These mutants secrete these proteins to the extracellular space via a pathway that is not mediated by EMVs. The prolonged lag phase and secretion of diverse proteins may be due to the intracellular damages such as protein misfolding and decrease of membrane integrity in Δ*hm1880* and Δ*hm2766*. Interestingly, EMVs from Δ*hm1880* and Δ*hm2766*, which have a prolonged lag phase, and Δ*hm3484* and Δ*hm3946*, which grew similarly to the parental strain, contained diverse protein bands other than P49 (Fig. 4). These results suggest that the mechanism of P49-selective protein transport to EMVs is impaired in these mutants. EMVs that are different from those of the parental strain may be produced in these mutants, such as those produced by explosive cell lysis (3). On the other hand, Δ*hm4090*, Δ*hm2418*, Δ*hm2192*, Δ*hm502*, Δ*hm2827*, Δ*hm2775*, Δ*hm2704*, and Δ*hm3986* selectively transported P49 to EMVs like the parental strain although their EMV productivity was increased or decreased compared with the parental strain. Detailed analyses of these strains such as transcriptomic and proteomic analyses are necessary to understand how each gene disruption affects EMV production.

### Conclusions

In this study, we established the screening method using a curvature-sensing peptide, nFAAV5-NBD, and successfully identified the genes related to EMV production by *S. vesiculosa* HM13 (Fig. 5). The findings indicated that protein quality control, extracellular environmental signal sensing, and glutamate metabolism are involved in EMV production. The targeted gene-disrupted mutants of Dcp (Δ*hm4090*), glutamate synthase (Δ*hm3484*), and Rhs-family protein (Δ*hm2827*) produced significantly larger amounts of EMVs than the parental strains (Fig. 3A) without growth defects (Fig. 3D). These strains harbored a major cargo protein P49 on their EMVs (Fig. 4). These strains are expected to be useful for the production of heterologous proteins as cargoes of EMVs by using P49 as a carrier. Further research is required to verify whether curvature-sensing peptide-based screening can be applied to other bacterial species and to elucidate in detail the functions of the genes identified in this study in EMV biogenesis.

**Fig. 5.**
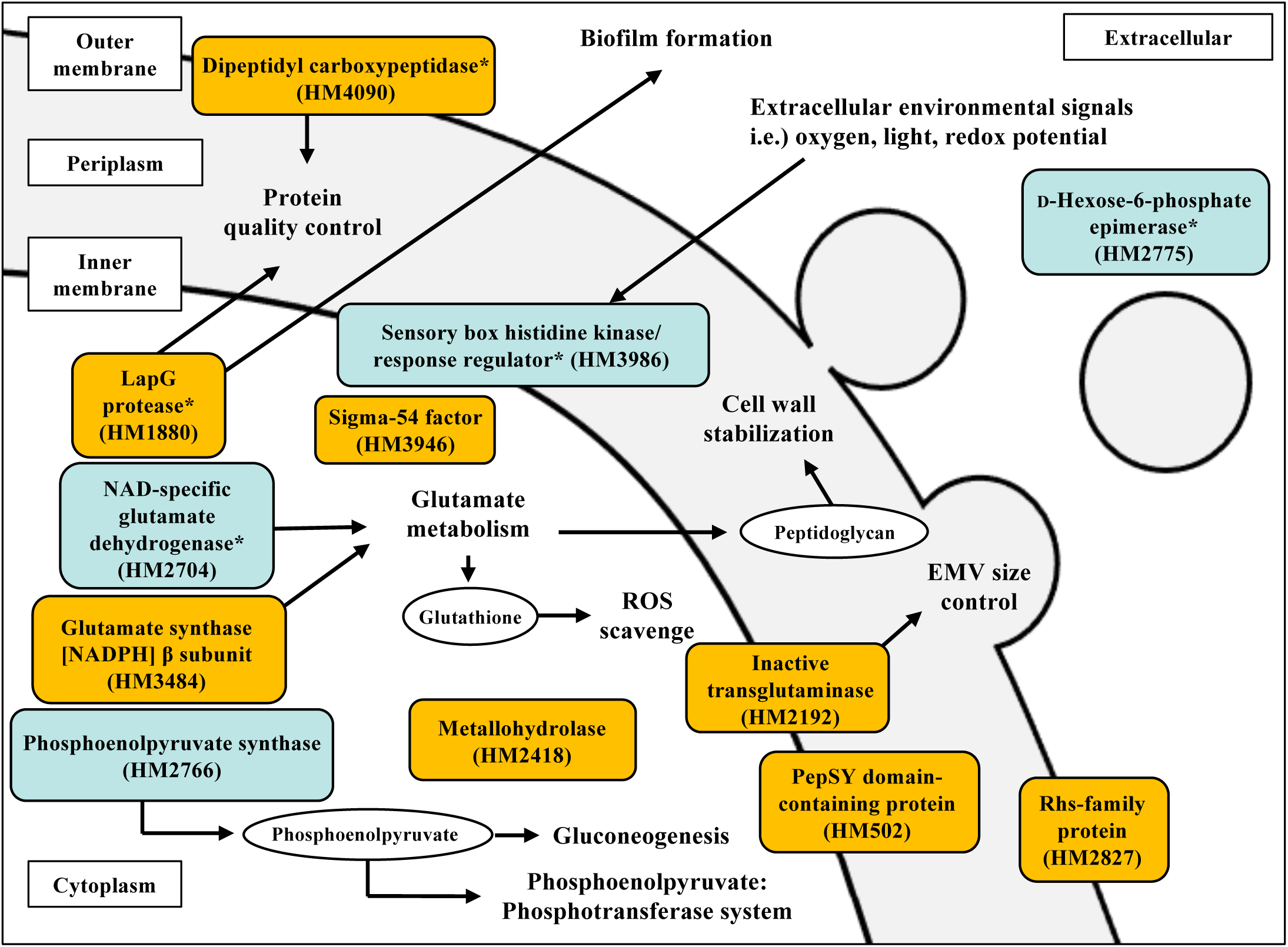
Proteins suggested to be involved in EMV production by *S. vesiculosa* HM13 in this study. Predicted localizations and functions of the proteins encoded by the genes whose disruptions increased (orange boxes) or decreased (blue boxes) EMV productivity are shown. Localizations were predicted by the PSORTb and SOSUI programs. Proteins with low localization reliability are marked with *.

## MATERIALS AND METHODS

### Bacterial strains and culture condition

The strains used in this study are listed in Table S3. The mutant of *S. vesiculosa* HM13, ΔPyrF^HM13^, with rifampicin (Rif) resistance and the gene of orotidine-5′-monophosphate decarboxylase (PyrF) deleted was used as the parent strain (45). *E. coli* S17-1/λ*pir* was used as the donor strain in the conjugal gene transfer (46). Gene disruption and random transposon mutagenesis were performed with a knock-out plasmid, pKNOCK-Km (23), and a transposon-containing plasmid, pMiniHimar RB1 (47), respectively (Table S3). ΔPyrF^HM13^ and its derivatives were grown at 18 °C in LB medium (1% of tryptone (Nacalai Tesque, Kyoto, Japan), 0.5% of yeast extract (Difco, Becton Dickinson, Franklin Lakes, NJ, USA), 1% of NaCl) (pH 7.0). *E. coli* S17-1/λ*pir* was cultivated at 37 °C in LB medium. When necessary, Rif, kanamycin (Km), and uracil (Ura) were added at 50 µg/mL. The cultivation was performed in BioShaker BR-43FL (Taitec, Saitama, Japan) at 180 rpm and evaluated with spectrophotometer UV-2450 (Shimadzu, Kyoto, Japan) by monitoring an optical density at 600 nm (OD600). The growth of ΔPyrF^HM13^ and its derivatives was evaluated by monitoring OD660 at 70 rpm and 18 °C (BioPhoto recorder TVS062CA, ADVANTEC Toyo, Tokyo, Japan).

### Isolation of EMVs by ultracentrifugation

ΔPyrF^HM13^ and its derivatives were grown in 5 mL of LB medium and harvested at the early stationary phase (OD600 of about 2.0). The culture was centrifuged at 6,800 × *g* for 10 min at 4 °C. The cells were washed with Dulbecco’s PBS (DPBS) (48) and resuspended in DPBS. The cell suspension was stored until use for the evaluation of nFAAV5-NBD binding specificity. The supernatant was centrifuged at 13,000 × *g* for 15 min at 4 °C to remove the remaining bacterial cells. The supernatant was filtered through a 0.45-µm pore polyethersulfone (PES) filter to remove the remaining debris, and the filtrate was collected as a cell-free supernatant fraction. The cell-free supernatant fraction was stored until use for the evaluation of nFAAV5-NBD binding specificity. EMVs were obtained by ultracentrifugation from 3 mL of filtrate at 100,000 × *g* (average centrifugal force) and 4 °C for 2 h with 50Ti rotor and Optimax centrifuge (Beckman Coulter, Brea, CA, United States). The supernatant after removing EMVs was collected as PVF. The pellets of EMVs were resuspended in 300 µL of DPBS and collected as an EMV fraction. PVF and the EMV fraction were used for the evaluation of nFAAV5-NBD binding specificity and the analysis of secretory proteins by SDS-PAGE and CBB staining.

### Evaluation of nFAAV5-NBD binding specificity

The EMV-containing fractions (culture, cell-free supernatant, and EMVs) and the EMV-free fraction (cell suspension and PVF) were diluted with LB medium (for culture, cell-free supernatant, and PVF) or DPBS (for EMVs and cell suspension). The concentration of each component in the original culture was defined as a relative concentration of 1.0. Each fraction (100 µL) was transferred to 96-well black flatbottom plates (Greiner, Frickenhausen, Germany), and then nFAAV5-NBD was added at a final concentration of 0.5 µM. After 5 minutes of incubation at room temperature, the fluorescence intensity of NBD was measured with Infinite^®^ 200 Pro M Nano+ (Tecan, Männedorf, Switzerland) at 480 nm excitation and 530 nm emission. The data were obtained under optimized experimental conditions (e.g., gain: 100, and the z-position of the luminescence fiber bundle: 20000 µm).

### Random transposon mutagenesis and screening using nFAAV5-NBD

The random transposon mutagenesis was conducted by conjugation of ΔPyrF^HM13^ and *E. coli* S17-1/λ*pir* harboring pMiniHimar RB1. Single colonies of Tn mutants formed on LB agar plates containing Rif, Km, and Ura were isolated and inoculated into 150 µL of LB liquid medium supplemented with Ura in 96-well culture plates. Following static culture for 24 h at 18 °C, all Tn mutants were subcultured in 150 µl of fresh LB media (1: 20 dilution rate) containing Ura and statically cultivated at 18 °C for 24 h when the OD600 of ΔPyrF^HM13^ reached about 1.8 (Fig. S4). After cultivation, 100 µL culture was transferred into 96-well black flatbottom plates (Greiner), and nFAAV5-NBD was added to each well at a final concentration of 0.5 µM. After 5 minutes of incubation at room temperature, the fluorescence intensity of NBD was measured with Infinite^®^ 200 Pro M Nano+ (Tecan) at 480 nm excitation and 530 nm emission. Fold change of EMV production was defined as follows: Fold change = [NBD fluorescence intensity/OD600]Tn/[NBD fluorescence intensity/OD600]parent (“Tn” and “parent” indicate the values of the Tn mutant culture and the parental strain culture, respectively). The strains showing more than 2.0- or less than 0.5-fold EMV production were selected as hyper- or hypo-vesiculating candidates, respectively (1st selection). The selected candidates were aerobically cultured in 5 ml of LB medium containing Ura (18 °C, 180 rpm (BioShaker BR-43FL, Taitec), 24 h), and EMV production was again evaluated using nFAAV5-NBD. Candidates that finally showed fold change of more than 1.6 and less than 0.8 were selected as hyper- and hypo-vesiculating strains, respectively (2nd selection).

### Identification of transposon insertion sites

To identify the insertion sites of the transposon, we performed rapid amplification of the regions adjacent to the transposon ends using a single primer (Table S4; Single-Primer-PCR-1, Single-Primer-PCR-2, or Single-Primer-PCR-3). All the Tn mutants were subjected to single-primer PCR (22), and the PCR products were sequenced with a nested primer (Table S4; Single-Primer-PCR-nested), resulting in identification of the transposon insertion sites of the Tn mutants except for strain 3-D3 and 18-G9. The genomic DNA extracts of strain 3-D3 and 18-G9 were digested with the restriction enzymes, XbaI (Takara Bio, Shiga, Japan) and SacI (Takara Bio), and circularized DNA prepared by self-ligation with Ligation high Ver.2 (Toyobo, Osaka, Japan) was used as a template for inverse PCR (21). The genomic DNA region flanked by the two ends of the transposon in the circular DNA was amplified and sequenced by two primers (Table S4; Himar1 and Himar615), which anneal specifically to different strands of the transposon. The amino acid sequences of the identified genes were subjected to BLASTP (https://blast.ncbi.nlm.nih.gov/Blast.cgi?PAGE=Proteins) and compared with protein sequences in the database. Proteins having sequence similarity and conserved domains are listed in Tables 1, 2, and S2. HM1880 and HM1528 did not have any specific sequence or domain hits in the database. Their sequences were submitted to AlphaFold2 version 1.4 (https://colab.research.google.com/github/sokrypton/ColabFold/blob/main/AlphaFold2.i pynb) to generate predicted conformational models. Based on these models, we searched for structurally similar proteins in Protein Data Bank (PDB) (https://www.rcsb.org/) using the Dali server (http://ekhidna.biocenter.helsinki.fi/dali/). Protein localization was predicted by PSORTb (https://www.psort.org/psortb/) and SOSUI programs (https://harrier.nagahama-i-bio.ac.jp/sosui/sosuigramn/sosuigramn_submit.html).

### Targeted gene disruption by single crossover recombination

A linear fragment of pKNOCK-Km was amplified with the primers pKNOCK-1 and -2 (Table S4). The internal fragments of each gene to be disrupted were amplified using the primers specific to the target genes (Table S4; hm4090-single-FW and -RV for *hm4090* disruption; hm3484-single-FW and -RV for *hm3484* disruption; hm1880-single-FW and -RV for *hm1880* disruption; hm2418-single-FW and -RV for *hm2418* disruption; hm2721-single-FW and -RV for *hm2721* disruption; hm3946-single-FW and -RV for *hm3946* disruption; hm2192-single-FW and -RV for *hm2192* disruption; hm502-single-FW and -RV for *hm502* disruption; hm3230-single-FW and -RV for *hm3230* disruption; hm2827-single-FW and -RV for *hm2827* disruption; hm2766-single-FW and -RV for *hm2766* disruption; hm2775-single-FW and -RV for *hm2775* disruption; hm369-single-FW and -RV for *hm369* disruption; hm2704-single-FW and -RV for *hm2704* disruption; hm3986-single-FW and -RV for *hm3986* disruption). The amplified linear fragment of pKNOCK-Km and the internal fragments of each gene were fused using In-Fusion^®^ HD Cloning Kit (Takara Bio, Shiga, Japan) according to the manufacturer’s instructions. The insertion of the internal fragments into an appropriate site on pKNOCK-Km was confirmed by PCR with a primer pair of pKNOCK-check-FW and pKNOCK-check-RV (Table S4). ΔPyrF^HM13^ was conjugated with *E. coli* S17-1/λ*pir* transformed with the plasmids, and then the recombinants were selected on 1.5% LB agar plates containing Rif, Km, and Ura. The insertion of plasmids into an appropriate site on the genome was confirmed by PCR with a primer pair of pKNOCK-check-RV and one of the following primers: hm4090-check-FW, hm3484-check-FW, hm1880-check-FW, hm2418-check-FW, hm2721-check-FW, hm3946-check-FW, hm2192-check-FW, hm502-check-FW, hm3230-check-FW, hm2827-check-FW, hm2766-check-FW, hm2775-check-FW, hm369-check-FW, hm2704-check-FW, and hm3986-check-FW (Table S4).

### Quantification of EMV productivity and particle size distribution analysis of EMVs

To compare EMV productivity and particle diameters between the strains, EMV fractions were subjected to NTA using NanoSight NS300 (Nanosight Ltd., Malvern, United Kingdom). EMV solution diluted with DPBS was injected into the apparatus and analyzed with a camera level of 16 and a detection threshold of 5. EMV productivity and size distribution were analyzed based on these data using the NTA3.4 Built3.4.4 software.

### SDS-PAGE and CBB staining

Each EMV fraction and PVF were mixed with 1/10 volume of 100% (w/v) trichloroacetic acid (TCA), incubated on ice for 15 min, and subsequently centrifuged at 20,300 × *g* for 15 min at 4 °C to precipitate proteins. The pellets were washed with ice-cold acetone by centrifugation at 20,300 × *g* for 5 min at 4 °C. The dried protein pellet was suspended in SDS sample buffer (0.063 M Tris-HCl, 5% (v/v) 2-mercaptoethanol, 2% (w/v) SDS, 5% (w/v) sucrose, 0.005% (w/v) bromophenol blue) and incubated at 95 °C for 5 min. The protein from 300 µL of culture was loaded onto 12.5% SDS-PAGE gel. After electrophoresis, the gel was stained with Coomassie Brilliant Blue G-250 (CBB) (Nacalai Tesque).

## ACKNOWLEDGEMENTS

This work was supported in part by JSPS KAKENHI (JP18K19178 and JP20K20570 for TK and JP20K05786 and JP23H02127 for JK), a Grant from the Institute for Fermentation, Osaka (L-2019-2-012 for TK), and ICR Grants for Promoting Integrated Research from Institute for Chemical Research, Kyoto University (for KK and TO). The authors would like to express their sincere gratitude to Prof. Shiroh Futaki at Institute for Chemical Research, Kyoto University for generously providing access to a NanoSight NS300 instrument.

