## Supplementary material for "Rapid screening and identification of genes involved in bacterial extracellular membrane vesicle production using a curvature-sensing peptide": Fig. S1

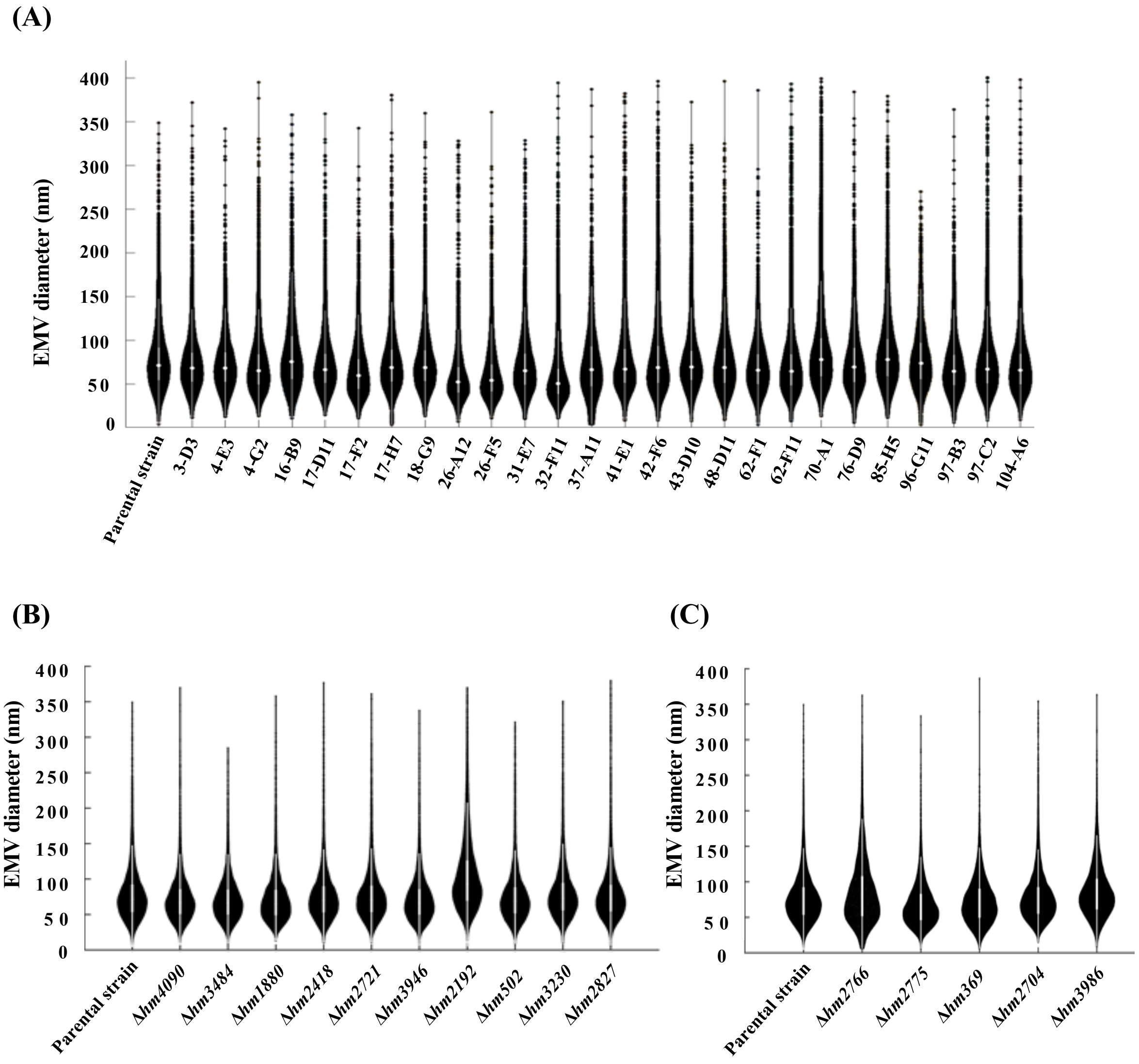


**Fig. S1 Violin plot of EMV hydrodynamic diameter**

The hydrodynamic diameter of EMVs was measured by NTA. (A) Transposon-inserted mutants. (B) Mutants obtained by disruption of the genes identified from hyper-vesiculating transposon-inserted mutants. (C) Mutants obtained by disruption of the genes identified from hypo-vesiculating transposon-inserted mutants.
