## Supplementary material for "Rapid screening and identification of genes involved in bacterial extracellular membrane vesicle production using a curvature-sensing peptide": Fig. S2

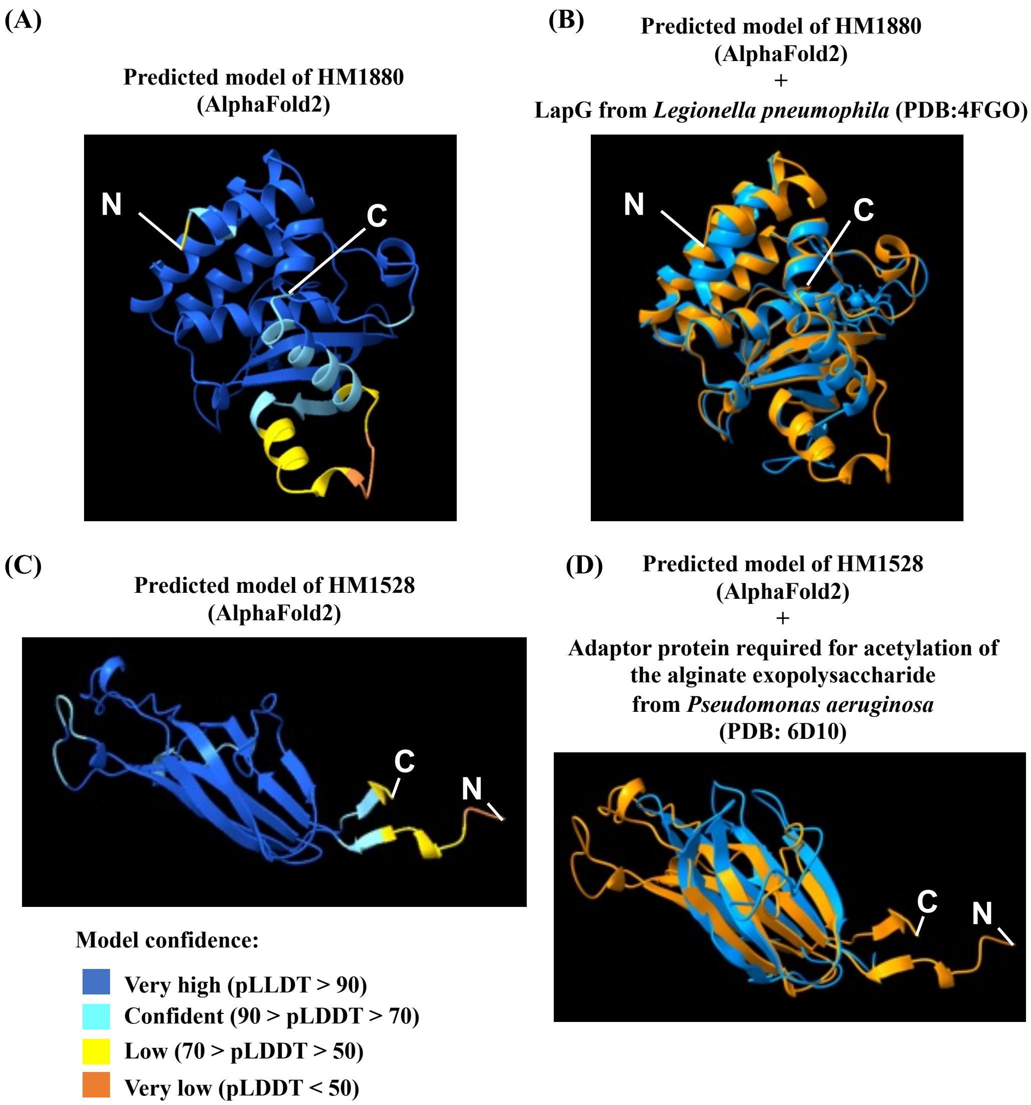


**Fig. S2 Prediction of structure for HM1880 and HM1528**

Conformational models of HM1880 without the N-terminal signal sequence (A) and HM1528 (C) were prepared by AlphaFold2. The residues were highlighted according to a per-residue confidence score (pLDDT). These predicted structures were overlaid with the protein conformations showing structural similarity to each protein (B and D). The orange structure models indicate HM1880 and HM1528, and the blue structures are the proteins acquired from PDB.
