## Supplementary material for "Rapid screening and identification of genes involved in bacterial extracellular membrane vesicle production using a curvature-sensing peptide": Fig. S3

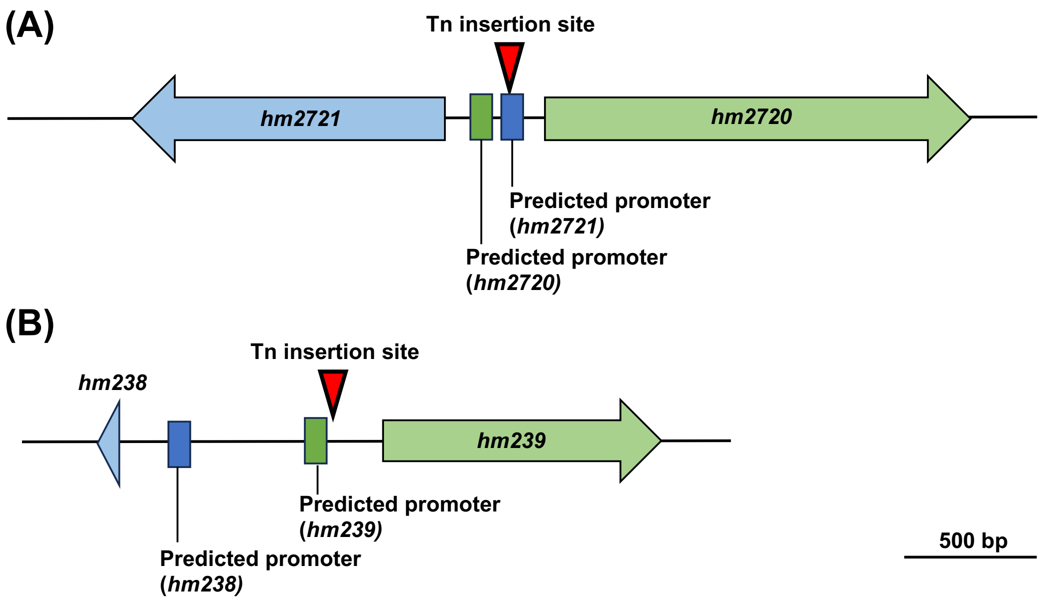


**Fig. S3 Transposon insertion sites of strain 26-F5 and strain 37-A11**
Transposon insertion sites were identified in strain 26-F5 (A) and strain 37-A11 (B). Promoter prediction was conducted using the neural network promoter prediction presented by Berkeley Drosophila Genome Project (https://www.fruitfly.org/seq_tools/promoter.html).
