## Supplementary material for "Rapid screening and identification of genes involved in bacterial extracellular membrane vesicle production using a curvature-sensing peptide": Fig. S4

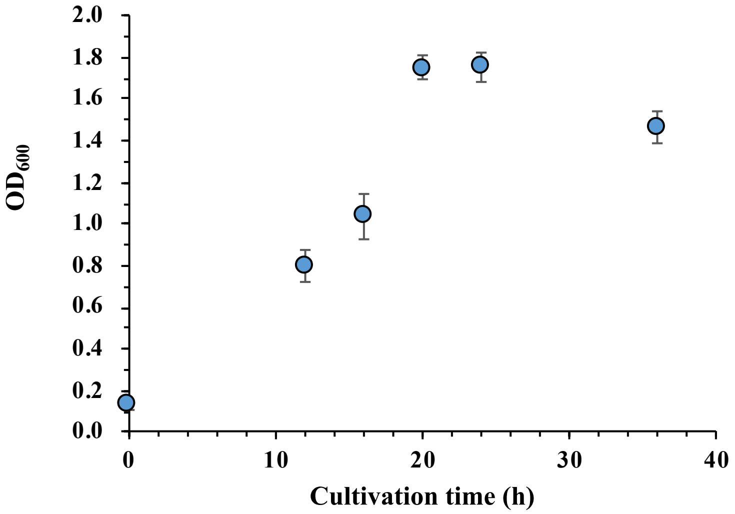


**Fig. S4 Growth of ΔPyrF$^{HM13}$ under the static cultivation condition in 96-well plates (Average +/- SD, n = 3)**

The growth of the parental strain under the static cultivation condition was examined. The strain grew to the stationary phase in 20−24 h of cultivation. In the 1$^{st}$ selection of Tn mutants, the bacterium was grown for 24 h to evaluate the EMV productivity of the mutants.
