## Supplementary material for "Rapid screening and identification of genes involved in bacterial extracellular membrane vesicle production using a curvature-sensing peptide": Table S1

### Table S1 Comparison of the relative EMV productivity evaluated by nFAAV5-NBD and NTA

| Hyper-vesiculating strain | 1st selection (nFAAV5-NBD) | 2nd selection (nFAAV5-NBD) | EMV productivity (NTA) |
|---|---|---|---|
| 3-D3 | 2.4 | 1.7 | 3.2 |
| 4-E3 | 2.1 | 1.7 | 2.4 |
| 4-G2 | 2.3 | 7.7 | 9.0 |
| 17-D11 | 3.1 | 1.6 | 4.4 |
| 17-F2 | 3.3 | 2.2 | 3.1 |
| 18-G9 | 2.4 | 2.5 | 2.9 |
| 26-F5 | 8.7 | 3.2 | 2.4 |
| 31-E7 | 2.8 | 15.7 | 1.7 |
| 32-F11 | 3.4 | 10.0 | 3.7 |
| 41-E1 | 2.3 | 4.1 | 1.2 |
| 42-F6 | 2.0 | 2.2 | 1.1 |
| 43-D10 | 2.5 | 3.3 | 1.5 |
| 62-F1 | 2.1 | 4.5 | 2.2 |
| 62-F11 | 2.3 | 7.2 | 6.1 |
| 70-A1 | 2.2 | 2.9 | 2.7 |
| 97-B3 | 7.6 | 2.8 | 1.9 |
| 97-C2 | 9.2 | 2.1 | 3.3 |
| 104-A6 | 3.4 | 2.5 | 1.7 |

| Hypo-vesiculating strain | 1st selection (nFAAV5-NBD) | 2nd selection (nFAAV5-NBD) | EMV productivity (NTA) |
|---|---|---|---|
| 16-B9 | 0.2 | 0.2 | 0.45 |
| 17-H7 | 0.5 | 0.8 | 0.33 |
| 26-A12 | 0.5 | 0.4 | 0.46 |
| 37-A11 | 0.4 | 0.6 | 0.41 |
| 48-D11 | 0.1 | 0.4 | 0.74 |
| 76-D9 | 0.4 | 0.3 | 0.44 |
| 85-H5 | 0.4 | 0.7 | 0.32 |
| 96-G11 | 0.5 | 0.5 | 0.31 |

The relative values are the EMV productivity of each mutant compared to the parent strain.
The first selection: $n = 2$, the second selection: $n = 3$, EMV productivity (NTA): $n = 3$.
