## Supplementary material for "Rapid screening and identification of genes involved in bacterial extracellular membrane vesicle production using a curvature-sensing peptide": Table S2

### Table S2 List of the domain hits searched by BLASTP

| Protein | Length (a.a.) | Name | Accession | Description | Interval | E-value |
|---|---|---|---|---|---|---|
| HM502 | 529 | PepSY_TM | pfam03929 | PepSY-associated TM region | 19–376 | 2.99E-60 |
| | | PiuB | COG3182 | Uncharacterized iron-regulated membrane protein | 19−381 | 2.71E-32 |
| HM1880 | 219 | COG3672 | COG3672 | Predicted transglutaminase-like cysteine proteinase | 54−192 | 4.85E-36 |
| | | Peptidase_C93 | pfam06035 | Bacterial transglutaminase-like cysteine proteinase BTLCP | 66−172 | 1.86E-11 |
| HM2418 | 363 | ComEC | COG2333 | Metal-dependent hydrolase, beta-lactamase superfamily II | 2−334 | 1.45E-11 |
| | | ComA-like_MBL-fold | cd07731 | Competence protein ComA, ComEC and related proteins | 4−270 | 2.37E-15 |
| | | Lactamase_B | smart00849 | Metallo-beta-lactamase superfamily | 31−98 | 1.44E-04 |
| HM2192 | 502 | Transglut_i_TM | pfam14400 | Inactive transglutaminase fused to 7 transmembrane helices | 25−184 | 7.40E-77 |
| | | 7TM_transglut | pfam14402 | 7 transmembrane helices usually fused to an inactive transglutaminase | 254−499 | 6.80E-133 |
| HM2704 | 1615 | Gdh2 | COG2902 | NAD-specific glutamate dehydrogenase | 26−1610 | 0.00E+00 |
| | | Bac_GDH | pfam05088 | Bacterial NAD-glutamate dehydrogenase | 76−1605 | 0.00E+00 |
| HM2766 | 808 | PRK06464 | PRK06464 | phosphoenolpyruvate synthase | 21−807 | 0.00E+00 |
| | | PEP_synth | TIGR01418 | phosphoenolpyruvate synthase | 23−806 | 0.00E+00 |
| | | PpsA | COG0574 | Phosphoenolpyruvate synthase/pyruvate phosphate dikinase | 23−806 | 0.00E+00 |
| | | PPDK_N | pfam01326 | Pyruvate phosphate dikinase, PEP/pyruvate binding domain | 35−369 | 8.46E-157 |
| HM2775 | 281 | YeaD | COG0676 | D-hexose-6-phosphate mutarotase | 6−272 | 2.90E-71 |
| | | D-hex-6-P-epi_like | cd09020 | D-hexose-6-phosphate epimerase-like | 16−278 | 7.86E-104 |
| | | Aldose_epim | pfam01263 | Aldose 1-epimerase | 19−278 | 2.02E-40 |
| HM2827 | 1557 | DUF6531 | pfam20148 | Domain of unknown function (DUF6531) | 156−241 | 7.76E-20 |
| | | RHS_repeat | pfam05593 | RHS proteins contain extended repeat regions | 469−502 | 2.13E-03 |
| | | YD_repeat_2x | TIGR01643 | YD repeat | 490−531 | 4.14E-03 |
| | | RHS | pfam03527 | RHS protein | 1207−1242 | 6.06E-12 |
| | | Rhs_assc_core | TIGR03696 | RHS repeat-associated core domain | 1260−1325 | 1.00E-33 |
| | | HopBF1 | cd20900 | Type III secretion system (T3SS) effector HopBF1 | 1369−1527 | 9.42E-22 |
| HM3484 | 468 | gltD | PRK12810 | Glutamate synthase subunit beta | 1−467 | 0.00E+00 |
| | | GltD | COG0493 | NADPH-dependent glutamate synthase beta chain or related oxidoreductase | 23−467 | 0.00E+00 |
| | | Fer4_20 | pfam14691 | Dihydroprymidine dehydrogenase domain II, 4Fe-4S cluster | 23−134 | 1.54E-56 |

| | | | | | | |
|---|---|---|---|---|---|---|
| HM3946 | 494 | PRK05932 | PRK05932 | RNA polymerase factor sigma-54 | 1−494 | 0.00E+00 |
| | | RpoN | COG1508 | DNA-directed RNA polymerase specialized sigma subunit, sigma54 homolog | 1−494 | 1.04E-180 |
| | | rpoN_sigma | TIGR02395 | RNA polymerase sigma-54 factor | 10−492 | 1.13E-164 |
| | | Sigma54_DBD | pfam04552 | Sigma-54, DNA binding domain | 335−492 | 1.23E-91 |
| HM3986 | 1236 | PAS_4 | pfam08448 | PAS fold | 465−573 | 3.40E-18 |
| | | PAS | COG2202 | PAS domain | 470−705 | 4.14E-21 |
| | | BaeS | COG0642 | Signal transduction histidine kinase | 670−953 | 9.23E-60 |
| | | PRK11107 | PRK11107 | hybrid sensory histidine kinase BarA; Provisional | 711−1228 | 1.35E-133 |
| | | HATPase_EvgS-ArcB-TorS-like | cd16922 | Histidine kinase-like ATPase domain of two-component sensor histidine kinases | 838−945 | 2.30E-55 |
| HM4090 | 716 | Dcp | COG0339 | Zn-dependent oligopeptidase | 44−716 | 0.00E+00 |
| | | M3A_DCP | cd06456 | Peptidase family M3, dipeptidyl carboxypeptidase (DCP) | 70−714 | 0.00E+00 |
| | | Peptidase_M3 | pfam01432 | Peptidase family M3 | 268−714 | 1.62E-149 |

The E-value is the number of hits expected with a similar score by chance when searching the database with the amino acid sequence of each protein.
