## Supplementary material for "Rapid screening and identification of genes involved in bacterial extracellular membrane vesicle production using a curvature-sensing peptide": Table S3

**Table S3 Bacterial strains and plasmids used in this study**

| Strains and plasmids | Descriptions | References |
|---|---|---|
| **Strains** | | |
| *Escherichia coli* S17-1/*λpir* | *E. coli* derivative, host for *pir*-dependent plasmids | (46) |
| $\Delta$PyrF$^{HM13}$ | Rifampicin resistant and *pyrF*-deletion mutant of *Shewanella vesiculosa* HM13 | (45) |
| Δ*hm4090* | *hm4090*-disrupted mutant (*hm4090*:pKNOCK-Km) of $\Delta$PyrF$^{HM13}$ | This work |
| Δ*hm3484* | *hm3484*-disrupted mutant (*hm3484*:pKNOCK) of $\Delta$PyrF$^{HM13}$ | This work |
| Δ*hm1880* | *hm1880*-disrupted mutant (*hm1880*:pKNOCK) of $\Delta$PyrF$^{HM13}$ | This work |
| Δ*hm2418* | *hm2418*-disrupted mutant (*hm2418*:pKNOCK) of $\Delta$PyrF$^{HM13}$ | This work |
| Δ*hm2721* | *hm2721*-disrupted mutant (*hm2721*:pKNOCK) of $\Delta$PyrF$^{HM13}$ | This work |
| Δ*hm3946* | *hm3946*-disrupted mutant (*hm3946*:pKNOCK) of $\Delta$PyrF$^{HM13}$ | This work |
| Δ*hm2192* | *hm2192*-disrupted mutant (*hm2192*:pKNOCK) of $\Delta$PyrF$^{HM13}$ | This work |
| Δ*hm502* | *hm502*-disrupted mutant (*hm502*:pKNOCK) of $\Delta$PyrF$^{HM13}$ | This work |
| Δ*hm3230* | *hm3230*-disrupted mutant (*hm3230*:pKNOCK) of $\Delta$PyrF$^{HM13}$ | This work |
| Δ*hm2827* | *hm2827*-disrupted mutant (*hm2827*:pKNOCK) of $\Delta$PyrF$^{HM13}$ | This work |
| Δ*hm2766* | *hm2766*-disrupted mutant (*hm2766*:pKNOCK) of $\Delta$PyrF$^{HM13}$ | This work |
| Δ*hm2775* | *hm2775*-disrupted mutant (*hm2775*:pKNOCK) of $\Delta$PyrF$^{HM13}$ | This work |
| Δ*hm369* | *hm369*-disrupted mutant (*hm369*:pKNOCK) of $\Delta$PyrF$^{HM13}$ | This work |
| Δ*hm2704* | *hm2704*-disrupted mutant (*hm2704*:pKNOCK) of $\Delta$PyrF$^{HM13}$ | This work |
| Δ*hm3986* | *hm3986*-disrupted mutant (*hm3986*:pKNOCK) of $\Delta$PyrF$^{HM13}$ | This work |
| **Plasmids** | | |
| pMiniHimar RB1 | Transposon insertion. Containing *pir* dependent R6K ori plasmid. $Km^r$. | (47) |
| pKNOCK-Km | Gene knock-out. Containing *pir* dependent R6K ori plasmid. $Km^r$. | (23) |
