## Supplementary material for "Rapid screening and identification of genes involved in bacterial extracellular membrane vesicle production using a curvature-sensing peptide": Table S4

**Table S4 Primers used in this study**

| Primers | Sequence (5'-3') | Target |
|---|---|---|
| pKNOCK-1 | GGGCTGCAGGAATTCGATATCAAGC | pKNOCK-Km |
| pKNOCK-2 | GGGGGATCCACTAGTTCTAGAGCG | As above |
| pKNOCK-check-FW | CCTCTCAAAGCAATTTTGAGTGACACAG | As above |
| pKNOCK-check-RV | TGCGTTTTCCCTTGTCCAGATAGC | As above |
| Himar1 | GGGAATCATTTGAAGGTTGGTAC | pMiniHimar RB1 |
| Himar615 | TTCTTCTGAGCGGGACTCTGGG | As above |
| Single-Primer-PCR-1 | TTCCAGTTTGAGATCTGG | As above |
| Single-Primer-PCR-2 | AAGAATAGACCGAGATAGG | As above |
| Single-Primer-PCR-3 | CCGAAATCGGCAAAATC | As above |
| Sigle-Primer-PCR-nested | CATTTAATACTAGCGACGCCATC | As above |
| hm4090-single-FW | ACTAGTGGATCCCCCTTTTTCAATCTGTCAGGGCTC | *hm4090* |
| hm4090-single-RV | GAATTCCTGCAGCCCGTTTCCCATAGCTGTTTACG | As above |
| hm4090-check-FW | CAACCGCTTATTTTTGAGTTTG | As above |
| hm3484-single-FW | ACTAGTGGATCCCCCTGTGCAAACAGTAGTGTATGACAAG | *hm3484* |
| hm3484-single-RV | GAATTCCTGCAGCCCCGCGAGCATTTTGTACTTCACG | As above |
| hm3484-check-FW | ATACATTACCGATACTGCCATATCACAAGG | As above |
| hm1880-single-FW | ACTAGTGGATCCCCCTTTAGCCAACCAGTTTGCTAAC | *hm1880* |
| hm1880-single-RV | GAATTCCTGCAGCCCCACGCACATACATCAAGCGC | As above |
| hm1880-check-FW | GCATGGATAAATATCGCCATTGGC | As above |
| hm2418-single-FW | ACTAGTGGATCCCCCCAGGGCTGACTGATAAGACAGG | *hm2418* |
| hm2418-single-RV | GAATTCCTGCAGCCCCCTGAATACCTGCATCCGCAG | As above |
| hm2418-check-FW | GGCCTGGAGGTGGTATTAAACG | As above |
| hm2721-single-FW | ACTAGTGGATCCCCCGTGGCTTGTTAGTGGCCAATG | *hm2721* |
| hm2721-single-RV | GAATTCCTGCAGCCCGCGCCAATGAGATAAACTCAACTG | As above |
| hm2721-check-FW | ATGCCATAGAACATATCCAAGGC | As above |
| hm3946-single-FW | ACTAGTGGATCCCCCGTGGCAAAAAAATCTCACGCC | *hm3946* |
| hm3946-single-RV | GAATTCCTGCAGCCCCTGGATTAAGGTGATTGCGTCACG | As above |
| hm3946-check-FW | CGACACAGCAGAATCCATGACC | As above |
| hm2192-single-FW | ACTAGTGGATCCCCCGCGACATCTATCGACATGATG | *hm2192* |
| hm2192-single-RV | GAATTCCTGCAGCCCAATAAGCTACCACCACCAGATAC | As above |
| hm2192-check-FW | TCGCCAACAACTGACATCATATG | As above |
| hm502-single-FW | ACTAGTGGATCCCCCCGAGCAAAAAAGGGCTGAATATCC | *hm502* |
| hm502-single-RV | GAATTCCTGCAGCCCATAACTGCCGCCAGGCAGATC | As above |
| hm502-check-FW | AGCCCATAGCTGGCTTGGAC | As above |
| hm3230-single-FW | ACTAGTGGATCCCCCTCCTGCCGGAGATGAGCTG | *hm3230* |
| hm3230-single-RV | GAATTCCTGCAGCCCTACGTCTTATGTCACCATGGAC | As above |

| | | |
|---|---|---|
| hm3230-check-FW | AAAACCGCGACCAACATGGC | As above |
| hm2827-single-FW | ACTAGTGGATCCCCCAACAAGATAATCAAGACAAGCCTG | *hm2827* |
| hm2827-single-RV | GAATTCCTGCAGCCCTTGATAAAAGTTCTGAGTATTCAGTAGC | As above |
| hm2827-check-FW | GGACCGCGACGGTGAATTAC | As above |
| hm2766-single-FW | ACTAGTGGATCCCCCGCTAAGCAAGCCATGGTGATTG | *hm2766* |
| hm2766-single-RV | GAATTCCTGCAGCCCTGTTTACCCGAGAAAATTAAGCCAGT | As above |
| hm2766-check-FW | ATCGGCAGTAAGTTGATTCAGATGG | As above |
| hm2775-single-FW | ACTAGTGGATCCCCCGCGATCAATCTGTTGCGTTGC | *hm2775* |
| hm2775-single-RV | GAATTCCTGCAGCCCAACCACATCGTCGACTTGTTC | As above |
| hm2775-check-FW | GGCGCAAATAGATCAATTTACACCC | As above |
| hm369-single-FW | ACTAGTGGATCCCCCAAACTCAAGCAAACCAAAACCAG | *hm369* |
| hm369-single-RV | GAATTCCTGCAGCCCTTTTCCTGGTTTAATGGCCAGTTTC | As above |
| hm369-check-FW | TAGTGGGTAGCTCTTTTGCAGGC | As above |
| hm2704-single-FW | ACTAGTGGATCCCCCCTTCTTACGCTCTGCCAAGTCA | *hm2704* |
| hm2704-single-RV | GAATTCCTGCAGCCCTACGAGTCTGATCTTTACAGTCTTG | As above |
| hm2704-check-FW | TTGTCTTGCCACGCTCAAGC | As above |
| hm3986-single-FW | ACTAGTGGATCCCCCGCAGAAAGAATGCTGGGTTACCG | *hm3986* |
| hm3986-single-RV | GAATTCCTGCAGCCCCTGAGAGCGATATTCCTGCACC | As above |
| hm3986-check-FW | TCGACTCAATCCCTGAAGCG | As above |

The sequences that anneal to pKNOCK-Km are underlined.
